# Cell expansion-division under resource limitation: a novel framework for modeling fruit growth dynamics

**DOI:** 10.1101/2024.05.30.596571

**Authors:** Leonardo Miele, Lionel Roques, Dario Constantinescu, Michel Génard, Nadia Bertin

## Abstract

Fruit growth is determined by cell expansion and division, processes that are closely linked to the availability and allocation of resources. By influencing environmental and growing conditions, agricultural practices affect these processes and ultimately many key indicators of fruit quality and yield. Given the multifaceted complexity underlying fruit growth dynamics, a process-based framework that can link genetic and environmental inputs to observables of economic and nutritional interest has become essential for predicting and improving agricultural outcomes. Here we present a mathematical framework that integrates cellular dynamics with resource allocation mechanisms to model the interplay between division, expansion and resource limitation in fruit cells. The model captures the temporal evolution of total fruit mass, cell number and cell mass distribution, reflecting data across genotypes and environmental conditions from tomato growth experiments. This sheds light on the possible relationships between genetic traits, growth conditions and fruit traits, and provides a predictive tool for optimizing fruit yield and quality under agricultural practices. Our minimal but mechanistic approach has broader implications for understanding resource-limited growth in multicellular systems.

## 1 INTRODUCTION

Fleshy fruits are the second most significant crop worldwide, due to their substantial economic, agronomic, and dietary contributions. Harvested fruits represent the culmination of complex processes involving cellular, multicellular and human components. Cell expansion determines the final size of the fruit (Musseau et al., 2017; Lemaire-Chamley et al., 2005), which is a crucial determinant for growers. Cell division is responsible for the production of differentiated tissue, which in turn determines the final total number of cells and their size distribution. This, in turn, affects a number of properties, including texture, firmness and nutrient content (Torres-Montilla and Rodriguez-Concepcion, 2021; Aurand et al., 2012). Both processes of expansion and division require carbon and nitrogen in order to function. These elements are transported via water and shared among cells, and are therefore subject to constraints that could limit the availability and allocation of resources.

Agricultural practices, by modifying the plant’s growing conditions, affect the environmental cues that regulate the cellular processes (Coen and Prusinkiewicz, 2024), resulting in considerable, genotype-dependent variations. For example, differences in fruit charge (*i*.*e*., the leaf-to-fruit ratio maintained on the plant) and irrigation regimes significantly influence fruit size as shape, as well as many organoleptic and nutrient properties (Bertin and Génard, 2018; Ripoll et al., 2016). While molecular and field experiments are essential for identifying and quantifying the relevant traits involved in fruit quality, process-based modeling aims to disentangle the complexity of such a Genotype×Environment×Practice (G×E×P) system (Fanta et al., 2014; Martre et al., 2011; Génard et al., 2007). By providing a mechanistic representation of the genetic, molecular and physiological processes at play, sound process-based approaches enable the connection between growth conditions and readily observable agronomic quantities (Baldazzi et al., 2012). This approach permits a parsimonious, *in silico* exploration of agricultural practices, thereby facilitating the formulation of predictions regarding improvements in fruit quality, yield indicators and water consumption (Baldazzi et al., 2019; Fanwoua et al., 2013).

Classical models of fruit growth often posit a temporal separation between periods of cell division and expansion. This assumption allows to capture the temporal patterns of either total cell number (Bertin et al., 2003) or total mass (Pantin et al., 2012; Liu et al., 2007; Génard et al., 2010). Similarly, models of resource limitation, which are used to explain mass growth patterns at various levels of plant organization (Génard et al., 2022; Prudent et al., 2014; Lescourret and Génard, 2003), are generally developed under the simplification of no cell division (Marcelis et al., 1998; Minchin et al., 1993). However, the expansion and division processes co-occur during the initial stages of fruit development (Renaudin et al., 2017; Pabón-Mora and Litt, 2011; Xiao et al., 2009), and their coordination is responsible for the typical sigmoid temporal patterns for mass and cell number, as well as for the long-tailed cell size distributions, ubiquitously observed in fruits (Bertin et al., 2002; Yamaguchi et al., 2002; Bain and Robertson, 1951) and leaves (Granier et al., 2000; Alves and Setter, 2004; Sunderland, 1960). Consequently, a framework aiming to simultaneously reproduce these three key observables must rely on a synergistic integration of expansion and division processes.

Expansion-division models have been extensively developed and applied to bacteria, plankton, microalgae and cancer cells (Miotto et al., 2024; Concas et al., 2016; Osella et al., 2014; Finkel et al., 2004). This research has focused on identifying the cell division mechanism responsible for the cell size distributions, resulting from *in vitro* exponential proliferation in the absence of nutrient, demographic or other constraints (Nieto et al., 2020). Instead, the dynamics of plant cells is markedly different from that of a clonal population exponentially colonizing a Petri dish, it cannot be explained by the simple sum of single-cell behaviors (Granier and Tardieu, 2009), and requires the invocation of both collective and environmental regulation arguments (Baldazzi et al., 2012).

In order to achieve this within a process-based framework, we propose a model which describes the dynamics of division and expansion within a population of cells which share a limited resource. Conceptually, our framework integrates the theory of size-dependent cell expansion-division processes (Perthame, 2006) with the source-sink formalism characterizing resource flow and sharing between cells (Allen et al., 2005; Thornley and Johnson, 1990). This latter argument is derived from typical plant physiology arguments, but its dynamical effect is expected to emerge in any context of resource limitation, regardless of its origin.

We apply this framework to both existing and novel empirical data, including experiments investigating the effects of genotype and of agricultural practices on tomato fruit growth. Tomatoes (*Solanum lycopersicum*) are a model species for fleshy fruit biology, due to their short life cycle, their great genotypic (Rothan et al., 2019) and phenotypic (Bergougnoux, 2014) diversity, and their economic and nutrient importance. Consequently, they represent a privileged system for the study of the effects of molecular mechanisms and agricultural practices on fruit growth (Mauxion et al., 2021). We successfully reproduce the temporal patterns observed in both total cell number and fruit mass. Furthermore, our model provides good agreement between predicted and observed cell mass distributions at harvest. We disentangle the different effects that genotype, fruit charge and water stress exert on the model parameters, providing valuable insights into their influences.

In conclusion, we discuss the implications of our findings for the optimization of fruit quantity and quality, and highlight potential extensions to further metabolic processes associated with cell size. By relying on minimal and general assumptions, our work also holds promise for application to other multicellular systems characterized by resource limitation.

## 2 MODEL DESCRIPTION AND DATA

The fruit is considered as a population of cells connected to the plant through resource pathways including xylem and phloem tissues (Thornley and Johnson, 1990). Acting as a source of resources, the plant supplies nutrients through the pathway to the fruit, driven by the sink force exerted by fruit cells. Fruit cells uptake these resources and convert them into new mass. Cells exist in two states: proliferating (*P*) and quiescent (*Q*). *P* cells both expand and divide, whereas *Q* cell can only expand. Furthermore *P* cells can also differentiate and turn into *Q*, but the reverse transition does not occur (Mauxion et al., 2021; Jones et al., 2019).

The central hypotheses of our model are the following: *i*) the processes of cell expansion and division depend on cell mass (Metz and Diekmann, 2014), and on the resources available (Thornley and Johnson, 1990); *i i*) the resource available to fruit cells is the result of a offer-demand flux balance between the source-plant and the fruit-sink (Minchin et al., 1993); *i i i*) cell differentiation from *P* to *Q* state occurs at constant rate.

The population’s state at time *t* is entirely determined by the densities (*i*.*e*. numbers per mass unit) *n*_*P*_ (*t, x*) and *n*_*Q*_ (*t, x*) of *P* and of *Q* cells, respectively, with mass *x*, which serves as a proxy for cell size. From these state variables, any macroscopic quantity of interest, such as the population total cell number *N* (*t*) or mass *M* (*t*), can be calculated through appropriate integration over the mass space, as summarized in Table 1. The aforementioned hypotheses are translated into mathematical formulas describing the mechanistic processes occurring at the single-cell level: an expansion function *ϕ* (*x*), describing the rate at which a cell of mass *x* increases its mass; a division function *γ* (*x*), describing the rate at which a parent cell of mass *x* divides into two offspring cells; a functional term Ω [*n* (*t*, ·)] describing resource-limitation, which depends on the state of the entire population (here, the *n* (*t*, ·) notation stresses the dependence on the state of the population across the mass space); a constant rate *ρ* at which *P* cells differentiate into *Q*.

**TABLE 1.** Notation of the state variables and functions.

| Symbol | Description |
| --- | --- |
| $x$ | Cell mass ($[\mu g]$) |
| $n_P(t, x)$ | Density of Proliferating cells with mass $x$ at time $t$ |
| $n_Q(t, x)$ | Density of Quiescent cells with mass $x$ at time $t$ |
| $n(t, x)$ | Total cell density $n_P(t, x) + n_Q(t, x)$ |
| $N(t)$ | Total Cell Number $\int_0^\infty n(t, x) dx$ |
| $M(t)$ | Total Fruit Mass $\int_0^\infty x n(t, x) dx$ |
| $\phi(x)$ | Single-cell expansion function |
| $\gamma(x)$ | Single-cell division function |
| $\Omega[n(t, \cdot)]$ | Population-level resource-limitation functional |

By invoking population balance laws on the above processes (Ramkrishna, 2000), the equations governing the temporal dynamics of the system are given by:

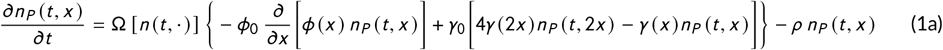

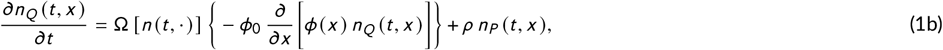

with:

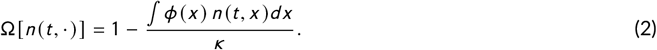

In Eq. (1a), the first term within the curly brackets represents the expansion process of *P* cells, equivalent to a transport in the mass domain with velocity *ϕ*. The parameter *ϕ*_0_ [*µg* ^1−[*ϕ*(*x*)]^ day^−1^] is a proportionality constant representing an effective expansion coefficient. This is the result of the compound contribution of several single-cell mechanistic parameters related to respiration cost, uptake rate, and mass conversion (details in the Appendix).

The second and third terms within the curly brackets represent the division process. The positive term refers to an increase in number due to the division of a parent cell of mass 2*x* into two progeny cells of equal mass *x*. In contrast, the negative term refers to a loss due to the dividing of the parent cells of mass *x* into two cells of half size. The parameter *γ*_0_ [day^−1^] represents the maximum rate at which division can occur. Estimating this value is complex, as the maximal division rate depends on numerous environmental conditions. To reduce the number of parameters, we adopt the estimate *γ*_0_ = 223.2 provided by Osella et al. (Osella et al., 2014). Using an alternative estimate would be tantamount to a rescaling of the two rate parameters involved in the model (*ϕ*_0_ and *ρ*); consequently, this would not alter our results (as they focus on shifts rather than on absolute values).

Finally, the last term represents the loss of *P* cells due to differentiation, which occurs at a constant rate *ρ [*day^−1^]. Similarly, in Eq. (1b), the term in curly brackets represents the expansion of *Q* cells, while the second term represents the gain due to *P* cells becoming *Q*. The division terms are absent because the *Q* cells are not subject to such a process.

The functional term Ω in Eq. (2) emerges from the source-sink balance between the plant and the fruit cells (details in the Appendix). It is a decreasing function of the total uptake demand coming from all cells (integral part), therefore it is expected to decrease over time as the fruit grows. Therefore, it is responsible for the deceleration of single-cell expansion over time, which subsequently affects all the other mass-dependent processes (in this case, solely division). The effective parameter *κ [µg* ^[*ϕ*(*x*)]+1^] is the result of the compound effect of both single-cell and plant and mechanisms, including uptake rate, phloem conductance and total resource concentration (details in the Appendix). In particular, in the limit of infinite resource concentration, one has *κ* → +∞, and hence Ω = 1 at any time. This implies that the total resource demand will never meet the plant offer and that growth is not constrained. In this sense, we interpret Ω as the contribution to population dynamics resulting from the sharing of the limited resource within cells, and *κ* as an effective resource limitation coefficient.

This formulation belongs to a novel class of Population Balance Equations (PBEs) accounting for population feedback due to resource limitation, where expansion and division events ultimately depend not only on single-cell state *x*, but also on the overall state of the population through Ω. Then, the standard PBE is recovered whenever Ω = 1, that is, when resource limitation can be neglected. The exact dynamics depends on the form of the expansion and division functions, whose specification can be driven by the knowledge of the biology of the system of interest. Due to its non-local, integro-differential nature, the system Eq. (1) cannot be solved analytical (Liou et al., 1997), but it has been integrated numerically upon adaptation of typical schemes. The numerical scheme proposed relies on choices and approximations required to overcome the complexities characterizing the systems’ dynamics. The scheme is detailed in the Supporting Information. The Python scripts can be found here.

### 2.1 Model functions and parametrization

According to standard considerations on cell physiology, cells grow via uptake of resource through the membrane. The uptake rate is considered proportional to the number of transporters present on the cell membrane, which in first approximation increases linearly with cell surface, leading to a 2/3-power of cell mass. Observations on meristem cells confirm this argument (R. Jones et al., 2017). Therefore, the mass-dependent expansion function is chosen as:

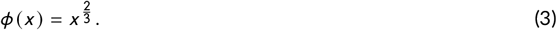

Contrary to prokaryotic cells, the absence of extensive real-time data on fruit cells does not allow us to infer the rate of division as function of mass. Instead of introducing further arbitrary, we use the parametric Hill function proposed in Osella et al. (2014):

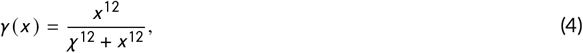

where the parameter *χ* [*µg*] determines the mass at which division rate reaches half-saturation.

In summary, we obtain a parametrization made of four parameters, one for each process (Table 2): *ϕ*_0_ for cell expansion, *χ* for cell division, *ρ* for cell differentiation, *κ* for resource limitation. The model is initialized with a genotype-dependent initial number of cells (all in the *P* state) in accordance with previous estimations (Baldazzi et al., 2019). The initial mass distribution is taken as peaked in small values, according to observations of tomato ovary cells (Renaudin et al., 2017; Musseau et al., 2017). Standard Bayesian parameter estimation is performed on the available datasets (details in the Appendix).

**TABLE 2.** Parameter definition and effects of agricultural practices on their best-fit values.

| Symbol | Description | Unit of measure | Water deficit | Fruit charge increase |
| --- | --- | --- | --- | --- |
| $\phi_0$ | Effective expansion coefficient | $\mu g^{1/3} \text{ day}^{-1}$ | ↘ | - |
| $\kappa$ | Resource limitation coefficient | $\mu g^{5/3}$ | ↘ | ↘ |
| $\chi$ | Half-saturation probability division size | $\mu g$ | ↗ | ↗ |
| $\rho$ | Differentiation rate | $\text{day}^{-1}$ | ↘ | ↘ |

### 2.2 Experimental datasets

The model is fitted on datasets from three different experiments (one already published and two original ones), conducted to investigate genotypic and environmental effects on tomato fruit growth. Details on Experiment 1 can be found in (Baldazzi et al., 2019), while for Experiments 2 and 3 we remand the Appendix.

In Experiment 1, tomato plants of a large-fruit - *Levovil* - and a cherry - *Cervil* - genotype are grown with either high or low fruit charge: plants cultivated with a high fruit charge are maintained with a low ratio between leaves and fruits (20 fruits per truss for *Cervil* and 6 for *Levovil*), whereas those with a low fruit charge exhibit a high leaf-to-fruit ratio (5 fruit per truss for *Cervil* and 2 for *Levovil*). The four datasets consist of the number of cells and the fruit mass measured at different stages of development, as well as the cell mass distribution measured on fruits at harvest.

In Experiment 2, cherry tomato plants, line *West Virginia 106* (*WVa106*) were grown in a glasshouse either under control or water deficit irrigation. In the control regime, water was supplied in order to meet the plant requirements based on the actual evapotranspiration. In the deficit irrigation regime, the water supply was reduced by half compared to the control. No cell mass distribution is provided for this experiment.

In Experiment 3, *Levovil* plants were grown in a glasshouse either under control irrigation (as above) or deficit irrigation induced by a 40% reduction in water supply. The two datasets consist of fruit mass measured at different stages as for the other experiments. However, here the number of cells is provided only at harvest (together with cell mass distribution).

## 3 RESULTS

### 3.1 Temporal patterns

Fig. 1 shows the temporal evolution of the total cell number and the mass throughout fruit growth. The solid curves correspond to the model’s predictions, obtained using the best-fit parameters. The scatter symbols correspond to the experimental data. Time is measured in Days After Anthesis (DAA).

**FIGURE 1.**
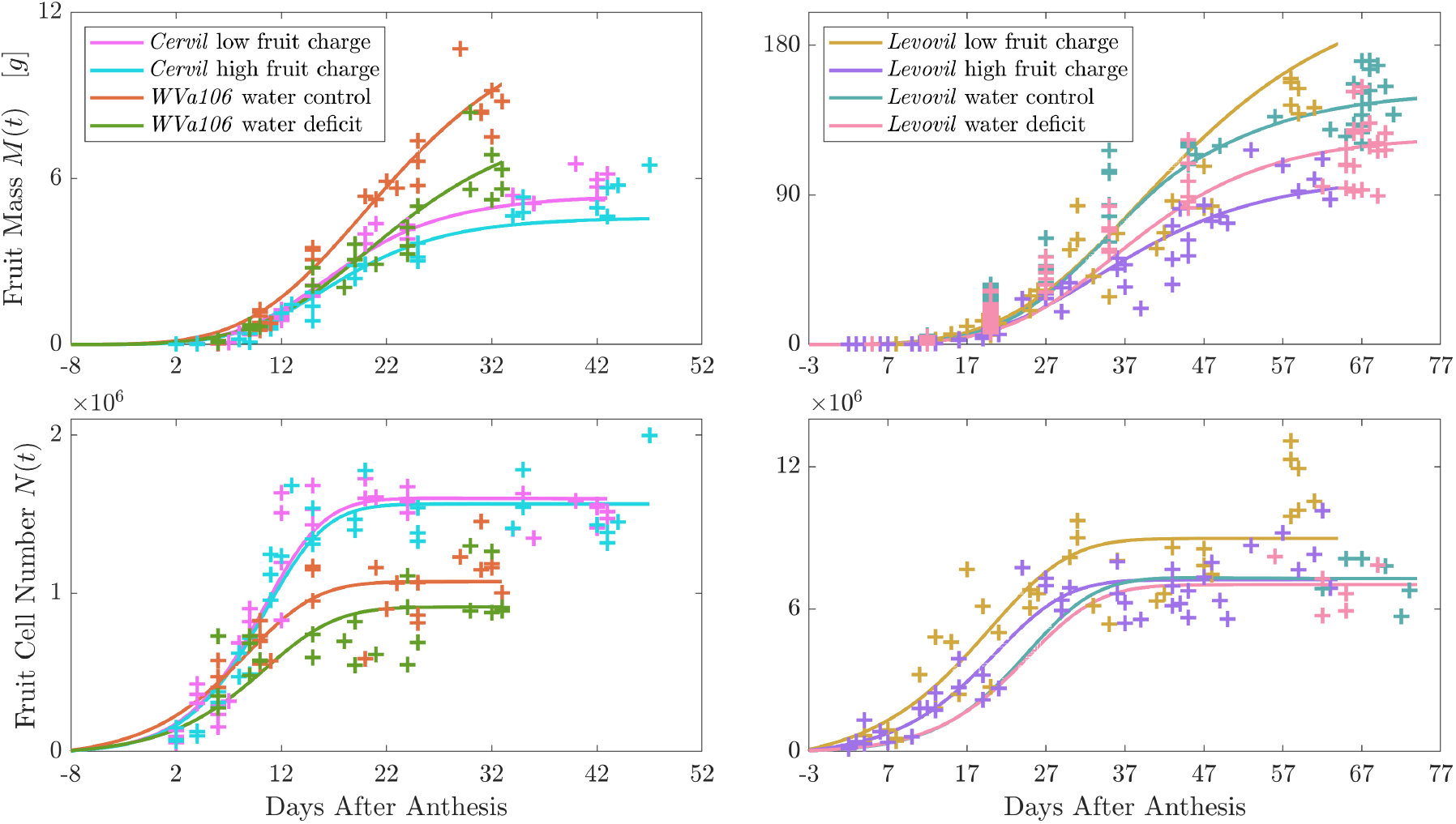
The temporal evolution of total fruit mass (top row) and total cell number (bottom row) is illustrated. The left column depicts small-fruit genotypes, while the right column depicts large-fruit genotypes. The color code, as specified in the legend, denotes the genotype and the agricultural practice involved. The solid lines represent the model’s output obtained with the best-fit parametrization, while the symbols represent the experimental data points.

The model is able to reproduce simultaneously the kinetics of cell number and fruit mass observed in all experiments. This encompasses the three-phase behavior that is characteristic of plant tissue. The initial phase is characterized by an exponential increase in both cell number and mass. In this phase, the population feedback Ω is approximately equal to one, indicating that the number kinetics is predominantly governed by cell division while cells expand at a rate approaching their maximal capacity. The duration of this phase ends at approximately DAA = 10 for cherry tomatoes, and at approximately DAA = 20 for large ones, in accordance with the experimental data.

As fruit mass increases, there is a concomitant decrease in cell resource uptake due to a reduction in the value of Ω, thereby initiating a second phase of linear growth. This phase exhibits slight asynchrony between numbers and masses due to the concurrent differentiation of *P* cells into *Q* cells, which contributes to the mitigation of division. The feedback between single-cell uptake and total fruit mass gives rise to a final, third phase. In this phase, the total cell number ceases to increase due to the decline in the cell division rate, which falls below the differentiation rate. This results in the emptying of the compartment of *P* cells.

Once all cells have entered a quiescent state, the total number reaches a plateau that determines the final number of cells at harvest. In contrast, fruit mass exhibits a sub-linear increase due to the approach of the feedback parameter to zero.

### 3.2 Cell mass distributions

Fig. 2 illustrates the temporal dynamics of the cell mass distribution for the *Cervil* low fruit charge case, obtained from a simulation of the model with best-fit parameters. The graph vividly illustrates a non-trivial behavior, characterized by a non-monotonic evolution of the function *n* (*t, x*). This is particularly evident at lower masses (the *x* axis is logarithmically scaled to emphasize this).

**FIGURE 2.**
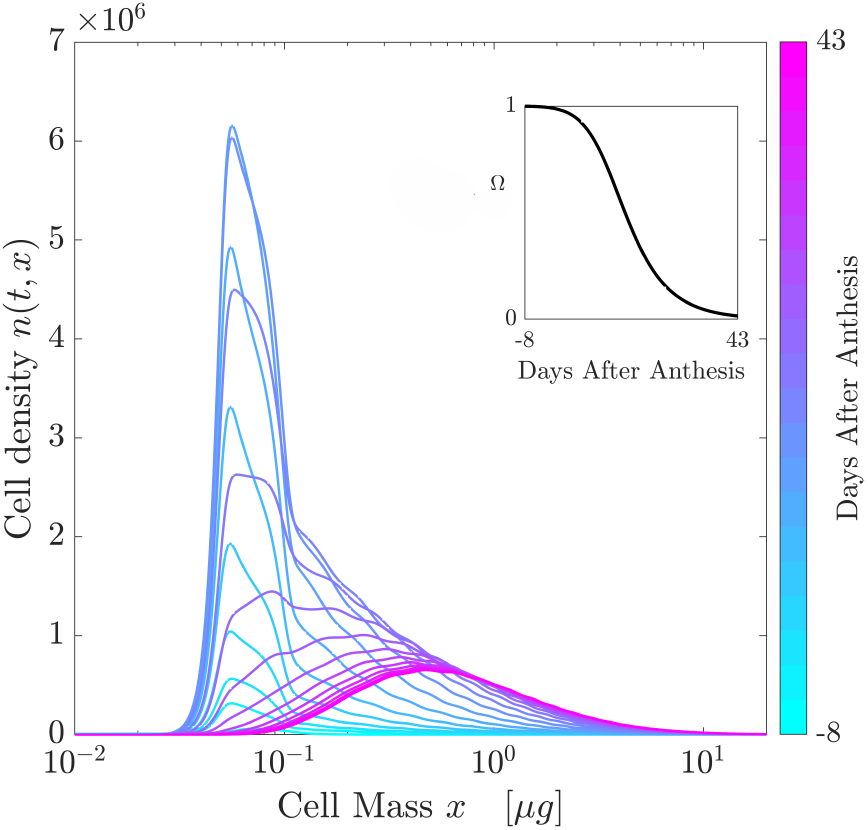
Dynamics of cell mass distribution in the *Cervil* low fruit charge dataset, reconstructed from numerical integration of Eq. (1) using the best-fit parameter values. The inset illustrates the kinetics of the population feedback contribution due to resource limitation, which causes the deceleration in the expansion-division processes.

Initially, the expansion process drives cells towards values of *x* close to half-saturation *χ*. The cells undergo extensive rounds of division around mass *χ*, continuously splitting into smaller cells which are promptly pushed by expansion toward values close to *χ*. This process gives rise to a cascade of division events that coincides with the initial exponential-like kinetic behavior. Concurrently, differentiation generates a subpopulation of cells that evade division and start an increase in mass above the value of *χ*, thereby contributing to cell heterogeneity.

Upon cessation of cell division due to extinction of the division rate, mass heterogeneity continues to increase, driven by differences in resource uptake provided by the mass-dependent function *ϕ* (*x*). The inset illustrates the temporal progression of the resource limitation contribution Ω, calculated numerically at each time step according to Eq. (2). The sigmoid shape is consistent with the aforementioned three-phase kinetics.

Fig. 3 compares the final distributions predicted by the model (obtained with the best-fit parameters) with the available experimental data measured at harvest. The model is able to reproduce the observed features, including the presence of non-trivial long tails and peaks in small mass values. It is important to note that these data were used exclusively for validation purposes and not for parameter estimation. In the absence of data for Experiment 2, the model provides the expected distribution.

**FIGURE 3.**
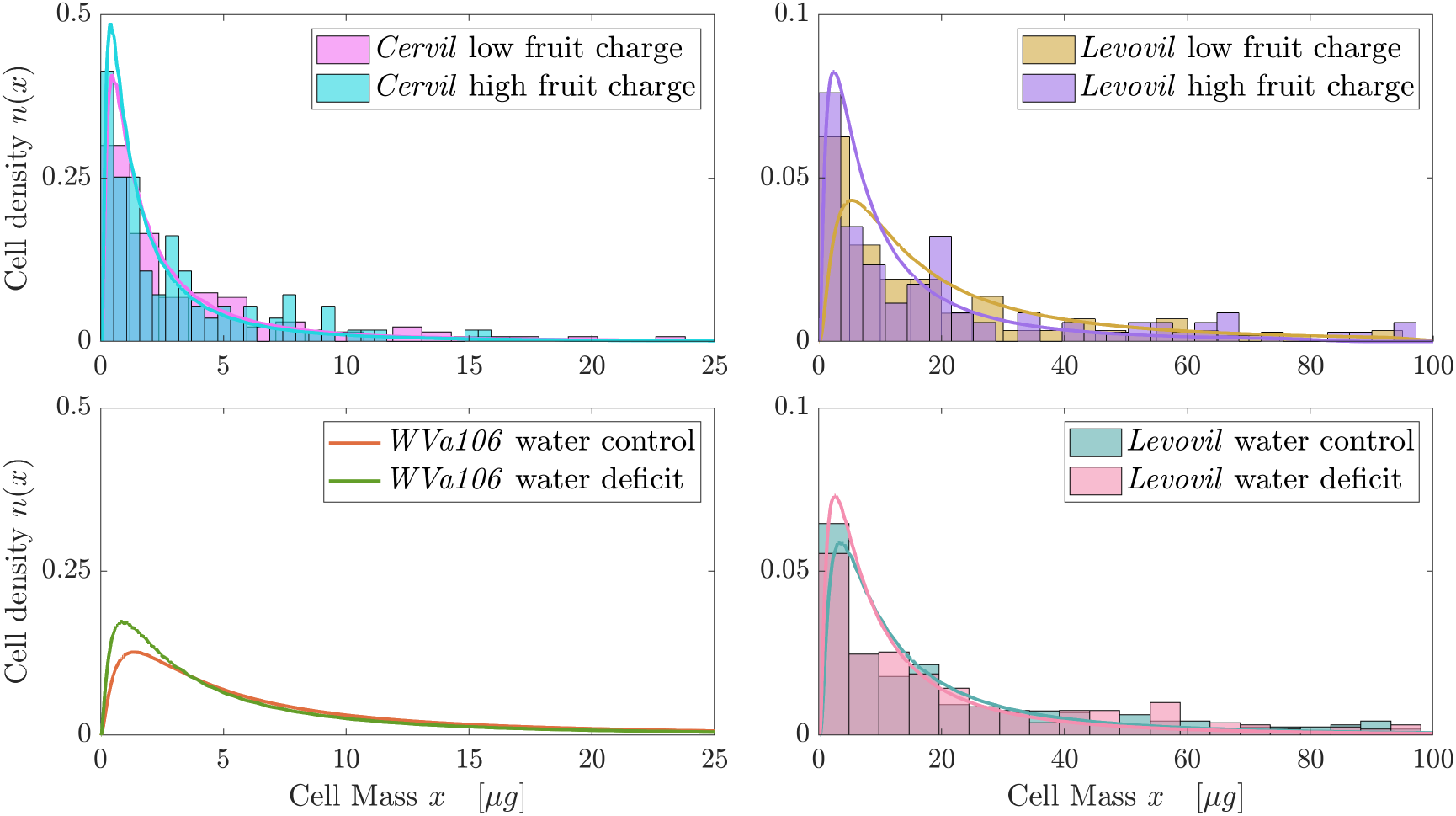
Cell mass density at harvest. The solid lines represent the prediction of the model with the best-fit parameters, while the histograms represent the empirical data.

### 3.3 Genotypic and environmental effects

The behavior of our G×E×P system is evaluated by examining how the values of the best-fit parameter change, when the genotype or agricultural practice changes. Our analysis cannot exclude the possibility that between-experiments differences may be due to different “absolute” conditions (*e*.*g*. greenhouse temperature, period of the year). Therefore, our focus is on within-experiment differences, that is, when different genotypes are grown at the same environmental conditions (Experiment 1), or when the same genotype is grown under different fruit charge or water regime (Experiments 1, 2 and 3). For the same reason, our interest is not on the absolute value of such estimations (summarised in Table S1 of the Supporting Information), but rather on statistically significant deviations (with respect to the Bayesian posterior distributions, shown in Fig. S1 of the Supporting Information).

The results are summarized in Table 2, showing increases (↗) or decreases (↘) of the best-fit parameter values with respect to agricultural practices. The growth conditions show minimal impact on the effective expansion coefficient *ϕ*_0_ across experiments. The differentiation rate *ρ* is significantly influenced by both the fruit charge and the irrigation conditions. The effective resource limitation coefficient *κ* is affected by both the genotype and the growth conditions. Large-fruit genotypes tend to exhibit higher value, whereas water-deficit and high fruit charge tend to reduce it. Finally, fruit charge affects the half-saturation mass *χ* in large-fruit genotypes but not in cherry ones.

## 4 DISCUSSION

Fruit growth depends on interrelated factors that remain incompletely understood. This is mainly due to the observational challenges presented by the complexity of such a G×E×P system. However, the significance of size-dependent behavior in plants is undeniable (Marshall et al., 2012), influencing both single-cell development and fruit quality traits. Here we have developed a process-based mathematical framework that focuses on minimal microscopic ingredients – expansion, division, differentiation and resource limitation – to explain and predict experimental patterns of fruit growth kinetics.

Our framework requires the estimation of only 4 parameters when applied to tomato fruit datasets, but is expected to work also on other single sigmoidal growth patterns, such as those observed in apples (Musse et al., 2021), cucumber (Marcelis and Baan Hofman-Eijer, 1993), olives (Camarero et al., 2024), eggplant, pepper and kiwifruit (Roch et al., 2020). The fitting procedure relies on macroscopic data (total cell number and mass) and is independent of single-cell mass distribution data. This offers several advantages. Firstly, it highlights the robustness of our approach and provides an autonomous validation of the chosen single-cell laws. Secondly, determining cell mass distributions in plant tissues involves sample disruption, rendering the process expensive and delicate, and variations in techniques may also introduce biases (McAtee et al., 2009).

By bypassing the necessity for cell mass distribution data in the fitting process, our model presents a cost-effective and non-invasive alternative to both make predictions on incomplete datasets (as we did for Experiment 2), and to perform *in silico* experiments (mimicking genetic or environmental modifications). Future applications of the model would benefit from precise estimations of cell numbers at very early stages. While evidence suggests environmental effects may manifest as early as the ovarian development period, their impact on pre-anthesis cell number appears limited (Aslani et al., 2023). Thus, such measurements could be leveraged to establish genotype-dependent initial conditions for the model, regardless of the growth conditions that will subsequently be employed during the experiment.

Contrary to previous works, our results reconcile cell- and mass-centered approaches, allowing for the reconstruction of both total cell number, fruit mass (Fig. 1) and cell mass distribution (Fig. 3). In addition, our model provides insights into the hidden dynamics of cell mass distribution during fruit growth (Fig. 2). The nonlinear trajectory shows that the interplay between cellular expansion, division, and nonlocal interactions gives rise to non-trivial dynamical patterns, which defy simple statistical models and reaffirm the advantage of dynamical ones. The emerging dynamics is consistent with temporal patterns observed experimentally in other fruit systems (Musse et al., 2021), and offers a mechanistic framework to support experimental research in food quality. For instance, cell mass distributions can be used to infer the micro-mechanical properties of cell walls (An et al., 2023), which in turn affect several fruit quality traits, such as: texture (Zdunek and Umeda, 2005), juiciness (Allan-Wojtas et al., 2003) and sugar content.

Instead of introducing temporal brakes that mimic metabolic regulation, our framework proposes resource limitation as the main mechanism responsible for the sigmoid kinetics typical of plant tissue: the balance between a finite resource supply and an increasing demand leads to the emergence of a contribution that depends on the whole population state (here represented by Ω). Ultimately, this contribution is responsible for the deceleration of the size-dependent processes (here expansion and division). Although we acknowledge the significance of metabolic regulation in coordinating the mechanisms that underlie plant cells development (Pinto et al., 2024; Jones et al., 2019), we argue that a shift towards a resource-based framework has several advantages: first, it lightens the parametrization and mitigates potential overfitting issues (Williamson et al., 2023; Apri et al., 2014); second, it facilitates a mechanistic understanding of the whole G×E×P system, as summarized in Table 2.

Increased fruit charge leads to smaller fruits, with fewer but larger cells on average. This results in a reduced resource availability (*κ*) and differentiation rate (*ρ*), and in an increased half-saturation division mass (*χ*). The decrease in *κ* can likely be attributed to phloem conductance corrections occurring at the fruit organizational level (see Eq. (10b)), consistent with previous findings (Génard et al., 2022). Interestingly, inspection of Eq. (10a) and Eq. (10b) suggests that potential genetic variations on single-cell resource uptake could be inferred by examining the inverse relationship between *κ* and *ϕ*_0_.

Some studies have proposed the existence of a compromise between resource uptake and fruit size, which would result in a trade-off between the values of *ϕ*_0_ and of *κ*. Our results do not permit a definitive conclusion to be drawn, as the estimated values of *ϕ*_0_ are found to be similar for both the large-fruit genotypes (*Levovil*) and the cherry genotypes (*Cervil* and *WVa106*). Nevertheless, our model could contribute to the validation of this trade-off, provided that further data on experiments where different genotypes are grown under the same environmental conditions becomes available.

The parameter *χ* is related to division: with larger values cells must be bigger to reach half the probability of division. Therefore, a larger value in conditions of lack of resource (high fruit charge or water deficit) suggests that in lack of resources cells tend to slower division. This could be a strategy of resource uptake, since cells exploit the resources to expand, rather than dividing. At the end of the process, indeed, the global number of cells is slower in lack of resources. Coherently, a lower value for the quiescence rate *ρ* in water deficit and high charge suggests that in lack of resources, cells tend to undergo more division events before entering into quiescence. While not clearly interpretable, this simulated phenomenon is interesting for studying the endoreduplication in further development of the model and deserves interest, since in many fruit species cells start an expansion process linked to ploidy increase (endoreduplication) when entering quiescence (Paige, 2018; Chevalier et al., 2011).

In this study, we have prioritized simplicity by assuming a constant resource supply throughout the dynamics. Nevertheless, our framework can be readily extended to incorporate more realistic, temporally varying agricultural practices. For instance, a common practice in processing tomato cultivation involves full irrigation during cell division and expansion, followed by a water deficit regime approximately 10 days before harvest. In our dynamical formalism, this approach would correspond to time-dependent functions for *κ* (*t*) and *ϕ*_0_ (*t*), which would modulate the expansion and division processes over time. These temporal variations would, in turn, influence fruit weight and composition, enabling direct comparisons with experimental data.

Future effort will be focused to the integration of metabolic modules of cell organelles production, such as plastids. Evidence show that the division rate of chloroplasts is subject to regulatory mechanisms related to both cell division and size (Chen et al., 2018). The abundance of chloroplasts is also considered to be a limiting factor for carotenoid formation, which are fundamental compounds for fruit organoleptic quality (Torres-Montilla and Rodriguez-Concepcion, 2021). Incorporating chloroplasts dynamics into our cell mass distribution evolution could therefore provide a tool to investigate chloroplast formations and its relationship with agricultural practices.

We conclude with a remark on the general significance of our study. We have obtained a novel class of PBEs that feature feedback between population and single-cell levels. This dynamics differs from that of previous PBE formulations, which considered extracellular resource dynamics (Henson, 2003), employed to model *in vitro* (Mantzaris et al., 1999) or bioreactor (Morchain et al., 2017; Concas et al., 2016) environments, where cell populations cease to proliferate due to resource depletion. We consider this mechanism to be inherent to all multicellular systems wherein cells draw resources from a shared, limited pool to fuel expansion and division. Therefore, we propose our formalism as a foundation for the analysis of the population dynamics of such systems. The derivation herein presented is based on fruit growth, but it is expected to be applicable to other plant tissues, such as leaves and seeds, as well as biofilm and cancer tissues, which exhibit similar behaviors (Vibishan et al., 2024; Bravo et al., 2023; Enrico Bena et al., 2021). The introduction of this new class of PBEs expands the range of dynamics related to cell expansion and division, paving the way for further applications and analysis.

## Abbreviations

G×E×P: Genotype×Environment×Practice;
PBE: Population Balance Equation
DAA: Days After Anthesis

## acknowledgements

The authors thank Gilles Vercambre and Valentina Baldazzi for fruitful discussions, Daniele Bevacqua for comments on an earlier version of the manuscript.

## 5 SUPPORTING INFORMATION

**S1**. PDE solver for the dynamical system

**S2**. Bayesian parameter estimation

**Table S1**. Best-fit parameter values and psoterior probability’s standard deviations.

**Figure S1**. Bayesian marginal posterior distributions of the parameters, for the *Cervil* and the *WVa106* datasets.

**Figure S2**. Bayesian marginal posterior distributions of the parameters, for the *Levovil* datasets.

## 6 APPENDIX

### 6.1 Microscopic processes - Cell expansion

In a minimal mechanistic description, a cell grows through uptake of the resource allocated to it. Part of such resource is converted into novel mass, while part of it is consumed in order to maintain the metabolic machinery. Cell expansion rate is proportional to a mass-dependent uptake function *ϕ* (*x*) (Génard et al., 2007), and to the concentration *R*_*f*_ of resource available to the fruit (generalization to non-linear cases is straightforward Génard et al. (2022), and is herein avoided in order to reduce the number of parameters). Therefore, a cell with mass *x* grows according to the following ordinary differential equation:

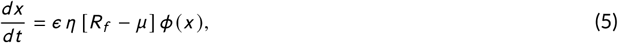

whe re *ϵ* [*µg*] is a resource-to-biomass conversion coefficient, *η* [day^−1^ *µg* ^−1−[*ϕ*(*x*)]^ *V*] is the single-cell uptake rate, *R*_*f*_ is the resource concentration available to the fruit, *µ* [µ*g V* ^−1^] is the metabolic discount, and *ϕ* (*x*) is the uptake per resource concentration function. Metabolic discount includes resource consumption by respiration, whose rate in tomatoes is proportional to mass growth rates (Grange and Andrews, 1995). The resource concentration *R*_*f*_ exploited by cells is obtained by invoking source-sink flux balance (Génard et al., 2022; Minchin et al., 1993; Thornley and Johnson, 1990). The resource flux ℱ [*µg* day^−1^] from the source to the sink is:

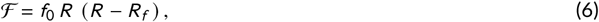

where *f*_0_[*µg* ^−1^ *V* ^2^day^−1^] is the transfer pathway’s conductance, and *R* [µ*gV* ^−1^] is the total resource concentration at the source. The unloading 𝒰[*µg* day^−1^] of resource at the sink is provided by the sum of all uptake contributions from all sink cells, that is:

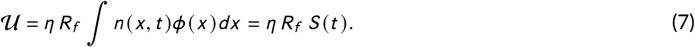

Assuming an absence of storage capacity within the transfer pathway, we have balance between in and out fluxes. Solving ℱ = 𝒰 for *R*_*f*_, we get:

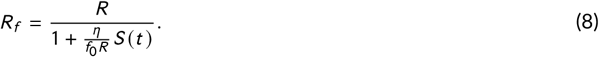

Inserting Eq. (8) into Eq. (5) we get:

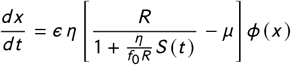

The three-parameter function 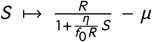 is convex. It equals *R* − *µ* at *S* = 0 and becomes zero at 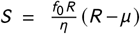.Thus, its shape mainly depends on the difference *R* − *µ* rather than solely on the values of *R* and *µ*. To avoid identifiability issues, we approximate this function with a simpler two-parameter linear function, which intersects the horizontal and vertical axes at the same points as the original function. This approximation leads to:

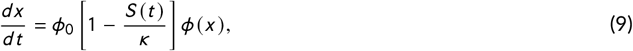

Where

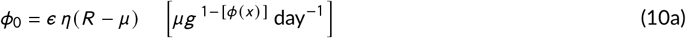

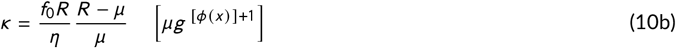

are the emerging effective compound parameters. Rather than on the five parameters *ϵ, η, f*_0_, *R* and *µ* separately, the model’s behavior is therefore best identifiable by the two effective parameters *ϕ*_0_ and *κ*, which can be fitted. Parameter *ϕ*_0_ increases proportionally to resource-mass conversion *ϵ*, single-cell uptake rate *η* and source resource concentration *R*, while it decreases proportionally to metabolic maintenance cost *µ*. Therefore, we refer to it as an effective single-cell expansion coefficient.

Parameter *κ* increases with phloem conductance *f*_0_ and total resource concentration *R*, while it decreases with single-cell uptake rate *η* and metabolic discount *µ*. We refer to *κ* as an effective resource limitation coefficient, as explained in the main text.

In order to lighten the notation employed in the main text, we write

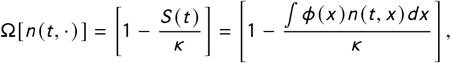

consistently with the terms appearing in Eq. (1).

#### 6.2 Microscopic processes - Cell division

The regulation of cell division is driven by both metabolic and environmental cues (Jones et al., 2019). In a mechanistic description of cell division, this complexity is generally captured by the introduction of size-dependent behavior. This so-called “sizer” mechanism represents a basic hypothesis of mathematical models (Facchetti et al., 2017) and is consistent with observations of root (Pavelescu et al., 2018) and meristem (R. Jones et al., 2017) cells. Accordingly, we define the probability density 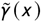 of a cell with mass *x* to divide. The corresponding division rate Γ, appearing in Eq. (1a), is obtained by following standard arguments (Liou et al., 1997). The probability of a cell to divide at a mass ∈ [*x, x* + *dx*] is 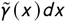. Let Δ*x* be the small mass increase in the small time interval Δ*t*, then the following relation holds:

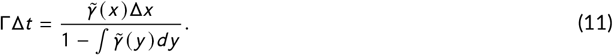

In the limit Δ*t* → 0, the single-cell expansion law Eq. (9) can be inserted in the above equation and the terms appearing in Eq. (1a) are obtained:

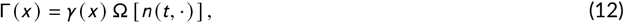

where 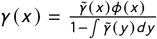 includes all the mass contributions to cell division (upon rescaling of the constant of proportionality with respect to single-cell uptake coefficient), while Ω [*n* (*t*, ·)] represents the same population contribution appearing in the mass expansion law.

#### 6.3 Experiment 2

In Experiment 2, cherry tomato plants (*Solanum lycopersicum*), line *West Virginia 106* (*WVa106*) were grown at a plant density of 2.2 plants *m* ^−2^ in a glasshouse in Avignon (south of France) from April to August. The day-night temperature set-point was 22^°^*C* − 16^°^*C*. Plants were planted in 7 *l* pots filled with potting soil (Klasmann, Substrat SP 15%) and an irrigation treatment was applied as soon as the first flowering truss appeared: 4 blocks of 5 plants were placed under “well-watered” irrigation conditions and 4 blocks of 5 plants were placed under “water-deficit” conditions, spread over 6 rows of 15 plants surrounded by border plants. The “well-watered” condition met the plant’s water requirements defined on the basis of the ETP measured in the greenhouse. For the water deficit, irrigation volumes were reduced by half compared with the control, without changing the frequency. The two treatments corresponded to a substrate water content of around 1.5 and 0.8 *g* H_2_O *g* ^−1^ dry soil for the control and deficit respectively. Nutrients were provided by fertigation (*EC* ≈ 2.0 to 3 *m S cm* ^−1^, *pH* ≈ 6) and no salinity stress occurred.

Flowering trusses were pruned to a maximum of 8 − 10 fruits, and flowers were hand-pollinated 3 times a week. Anthesis was recorded on each truss and fruits were harvested according to the number of DAA. Fruits were harvested between 2 DAA and maturity on 5 different trusses avoiding distal positions in the inflorescence. Fresh and dry fruit weights were measured as soon after sampling and after drying in a ventilated oven at 65^°^*C*. The number of pericarp cells was measured after enzymatic digestion of the walls and release of suspended cells as described in (Bertin et al., 2003).

Cells were counted in aliquots of a cell suspension under an optical microscope using Fuchs–Rosenthal chambers and Bürker chambers for the large and small fruit, respectively. Six to eight aliquots per fruit were observed and the cell number for the whole pericarp was calculated according to the dilution and observation volumes. In addition, cell size distributions (smallest and largest radii and 2D-surface) in the cell suspension aliquots were measured on ripe fruit using the ImageJ software (https://imagej.nih.gov/ij/). Surfaces are converted into masses by assuming that the cell is an ellipse made of water. Five samples of ∼ 20–25 cells per fruit pericarp were measured randomly for different fruits, so that around 300 − 500 cells were counted for each treatment.

#### 6.4 Experiment 3

*Levovil* plants (*Solanum lycopersicum*) were grown at a plant density of 2.4 plants *m* ^−2^ in a glasshouse in Bellegarde (South of France) from 1st February to June. The day-night temperature regime was monitored to 20 − 16^°^*C*. From flowering of the third truss, two water regimes were applied: well-watered plants were irrigated according to current practices (25% drainage); water-deficit plants were irrigated with a 40% decrease of water supply. Nutrients were provided by fertigation (*EC* ≈ 2.0 to 3 *m S cm* ^−1^, *pH* ≈ 6) and no salinity stress occurred. Flowering trusses were pruned to a maximum of 6 fruits, and flowers were pollinated by bumblebees.

Fruits were harvested between 5 DAA and maturity (red-ripe stage) on 5 different trusses avoiding distal positions in the inflorescence. Fresh and dry fruit weights were measured as soon after sampling and after drying in a ventilated oven at 65^°^*C*. At the red ripe stage, pericarp cell number and cell size distributions were measured on 6 fruits of each treatment, as described for Experiment 2.

