## Supplementary Information for "Cell expansion-division under resource limitation: a novel framework for modeling fruit growth dynamics"

### 1 PDE solver for the dynamical system

In addressing the challenges presented by our model — specifically, initial conditions concentrated near 0 with rapid decay, the need for precision in capturing phenomena close to 0 to accurately describe early cell divisions, ensuring total population conservation in the absence of a reaction term, minimizing numerical diffusion to preserve the distinct effects of growth and division, and the complexity introduced by nonlocal terms that complicate the use of implicit or Crank-Nicolson schemes — we embarked on a comprehensive evaluation of numerical methods. Initial tests with upwind and Lax-Friedrichs schemes revealed excessive numerical diffusion. Although the Lax-Wendroff scheme, enhanced with flux limiters, showed promise in simpler scenarios by significantly reducing numerical diffusion, its application to our problem was hindered. The scheme's reliance on the assumption that time and space derivatives are proportional by a constant factor proved inadequate for our complex model, where such a simplification does not hold, and any attempt to adapt it would require extensive modification.

To overcome these challenges and limitations, we developed a numerical scheme that balances the need for precision, mass conservation, and minimal numerical diffusion. The numerical method uses an explicit Euler scheme in time, a non-uniform grid in space, with grid points more concentrated for small $x$ values. The method deals separately with the transport and reaction terms in two separate steps. To deal with the transport equation, we first construct a spatial interpolation of the solution at time $t$ over $(0, L)$; secondly, we compute the exact solution at time $t + dt$ based on the solution at time $t$. To deal with the reaction term, we directly compute the terms on the right-hand side and use an interpolation to compute the values at positions $2x$. A commented Python version of the solver is available as a notebook here.

### 2 Bayesian parameter estimation

Mechanistic-statistical modeling bridges the gap between the solutions of continuous state models, such as our dynamical model, and empirical data, including noisy discrete data. This approach combines a mechanistic model, a probabilistic observation model that conditionally describes the data based on model outputs, and a statistical inference procedure. Initially applied to physical models and data [1], mechanistic-statistical methods have become increasingly standard in spatial ecology [2]. These methods require a mechanistic model that is not overparameterized, which may explain why their application at the organismal scale, such as in the case of fruits, remains rare.

#### 2.1 Observation model

The datasets are associated with the solution of Eq. 1 of the main text by positing that, for each dataset, the observed number of cells $\hat{N}(t)$ and the observed fruit mass $\hat{M}(t)$ at time $t$ adhere to Normal distributions. These distributions have mean values $N(t)$ and $M(t)$ (as defined in Table 1 of the main text), and standard deviations $\sigma_N$ and $\sigma_M \times M(t)$, respectively. The latter assumption is grounded in the observation that the standard deviation of the mass measurements tends to be proportional to the mass itself. For each dataset, $\sigma_N$ and $\sigma_M$ are calculated directly as the average standard deviation in the data (and the standard deviation divided by mass, respectively) between replicate observations (i.e. observations made on the same date).

#### 2.2 Statistical inference

The parameters $\phi_0$, $\kappa$, $\chi$, and $\rho$ are unknown. We employed a Bayesian method [3] to estimate their posterior distribution. The likelihood $\mathcal{L}$ represents the probability of observations given the parameters. Using the observation model and assuming independence of observations conditionally on the underlying biological process, we define:

$$\mathcal{L}_{\mathcal{N}}(\phi_0, \kappa, \chi, \rho) := P(\text{dataset}|\phi_0, \kappa, \chi, \rho) = \prod_{t \in \mathcal{T}_N} \prod_{i \in \#_t} \exp\left(-\frac{|\hat{N}_i(t) - N(t)|^2}{2\sigma_N^2}\right),$$

for the likelihood associated with the number of cells, and:

$$\mathcal{L}_{\mathcal{M}}(\phi_0, \kappa, \chi, \rho) := P(\text{dataset}|\phi_0, \kappa, \chi, \rho) = \prod_{t \in \mathcal{T}_M} \prod_{i \in \#_t} \exp\left(-\frac{|\hat{M}_i(t) - M(t)|^2}{2\sigma_M^2\, M(t)^2}\right),$$

for the likelihood associated with fruit mass. Here, $\mathcal{T}_N$ and $\mathcal{T}_M$ correspond to the measurement times and $\#_t$ to the corresponding number of replicate observations. The overall likelihood $\mathcal{L}(\phi_0, \kappa, \chi, \rho) = \mathcal{L}_{\mathcal{N}} \times \mathcal{L}_{\mathcal{M}}$ depends on the unknown parameters through $N(t)$ and $M(t)$.

The posterior distribution reflects the parameter distribution conditional on the observations:

$$P(\phi_0, \kappa, \chi, \rho|\text{dataset}) = \frac{\mathcal{L}(\phi_0, \kappa, \chi, \rho)\, \pi(\phi_0, \kappa, \chi, \rho)}{C},$$

where $\pi(\phi_0, \kappa, \chi, \rho)$ denotes the prior distribution of the parameters (as detailed below) and $C$ is a normalization constant independent of the parameters.

Finally, for the prior distributions, we adopted independent, non-informative uniform prior distributions for all parameters, with $\phi_0 \in (0, 1)$, $\kappa \in (5 \cdot 10^4, 2 \cdot 10^9)$, $\chi \in (0.05, 1)$, and $\rho \in (0, 1)$.

The numerical computation of the posterior distributions was performed using a Metropolis-Hastings (MCMC) algorithm with at least $2 \cdot 10^5$ iterations. The mean CPU time for solving Eq. 1 of the main text was approximately 25 $s$, with the total CPU time for the eight datasets being about 16 months (conducted in parallel for 2 months, on Intel(R) Xeon(R) CPU E5-2637 v3 @ 3.50GH).

The corresponding marginal distributions of $\phi_0$, $\kappa$, $\chi$ and $\rho$ are shown in Figs. 1-2. We note that these distributions are far from flat, indicating that the datasets do contain information on the parameters. The best-fit values are highlighted with circle symbols, while the error bars represent the standard deviations of the corresponding distributions (values also in Table 1). In constructing Table 2 of the main text, we considered shifts in the value of the best-fit parameters to be statistically significant if the error bars did not overlap.

Table 1: Best-fit parameter values and posterior probability's standard deviations

| Genotype | Treatment | $\phi_0$ | $\kappa$ | $\chi$ | $\rho$ |
|---|---|---|---|---|---|
| *Cervil* | low fruit charge | 0.34 ± 0.01 | 9.2e6 ± 3.2e5 | 0.60 ± 0.01 | 0.12 ± 0.01 |
| *Cervil* | high fruit charge | 0.33 ±0.01 | 8.5e6 ± 2.8e5 | 0.56 ± 0.01 | 0.12 ± 0.01 |
| *WVa106* | water control | 0.35 ± 0.02 | 1.4e7 ± 1.7e6 | 0.88 ± 0.02 | 0.07 ± 0.01 |
| *Wva106* | water deficit | 0.35 ± 0.01 | 1.0e7 ± 8.7e6 | 0.72 ± 0.01 | 0.11 ± 0.01 |
| *Levovil* | low fruit charge | 0.29 ± 0.01 | 2.4e8 ± 2.9e7 | 0.68 ± 0.02 | 0.09 ± 0.01 |
| *Levovil* | high fruit charge | 0.31 ± 0.02 | 1.2e8 ±2.1$e$7 | 0.55 ± 0.03 | 0.16 ± 0.03 |
| *Levovil* | water control | 0.40 ± 0.004 | 1.3e8 ± 4.7 e6 | 0.48 ± 0.01 | 0.44 ± 0.01 |
| *Levovil* | water deficit | 0.36 ± 0.004 | 1.2e8 ± 4.9e6 | 0.46 ± 0.01 | 0.37 ± 0.01 |

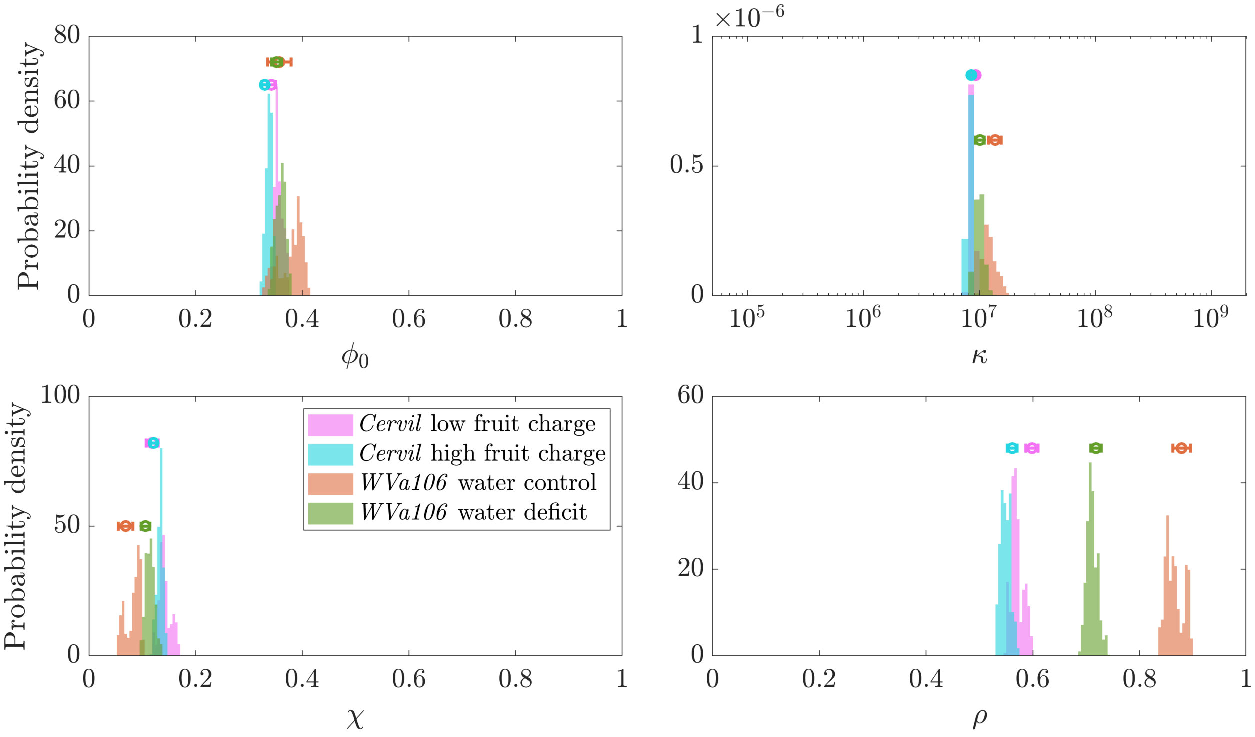


Figure 1: Bayesian marginal posterior distributions of the parameters, for the *Cervil* and *WVa106* datasets.

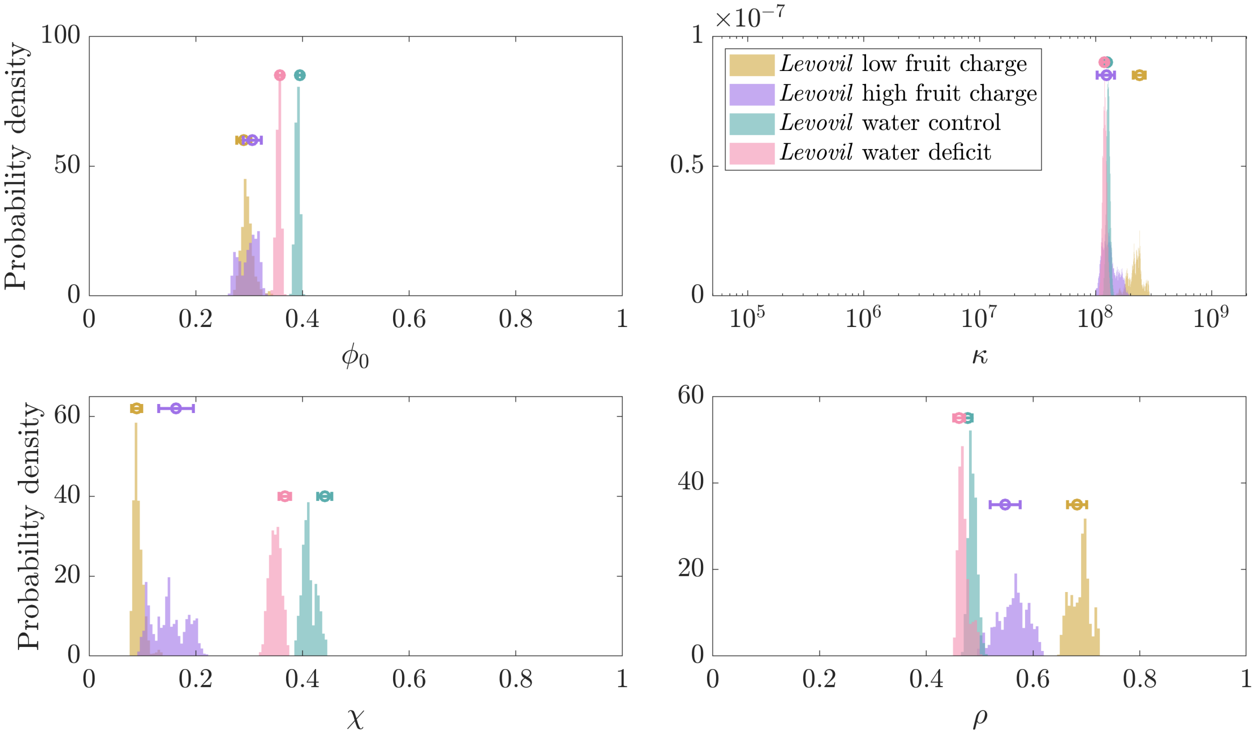


Figure 2: Bayesian marginal posterior distributions of the parameters, for the *Levovil* datasets.
